# Detecting mtDNA effects with an Extended Pedigree Model: An Analysis of Statistical Power and Estimation Bias

**DOI:** 10.1101/2024.12.19.629449

**Authors:** Xuanyu Lyu, S. Alexandra Burt, Michael D. Hunter, Rachel Good, Sarah L. Carroll, S. Mason Garrison

**Author notes:** Correspondence may be addressed to Xuanyu Lyu via.

## Abstract

Mitochondrial DNA (mtDNA) plays a crucial role in numerous cellular processes, yet its impact on human behavior remains underexplored. The current paper proposes a novel covariance structure model with seven parameters to specifically isolate and quantify mtDNA effects on human behavior. This approach uses extended pedigrees to obtain estimates of mtDNA variance while controlling for other genetic and environmental influences. Our Monte-Carlo simulations indicate that a sample size of approximately 5,000 individuals is sufficient to detect medium mtDNA effects (*mt*^2^ = 5%), while a more substantial cohort of around 30,000 is required for small effects (*mt*^2^ = 1%). We show that deeper pedigrees increase power to detect the mtDNA effect while wider pedigrees decrease power, given the equal total sample size. We evaluated how missing kinship records and mtDNA mutations impact bias. Both lead to underestimation of mtDNA variance, and an overestimation of the interaction between nuclear DNA and mtDNA. In addition, the false positive rate of mtDNA effect estimation is low when fitting the model with data generated without mtDNA effects. Collectively, we demonstrate that using extended pedigrees to quantify the influence of mtDNA on human behavior is robust and powerful.

## Introduction

### MtDNA Effect Estimation in Extended Pedigrees

Mitochondrial DNA (mtDNA) has garnered significant attention due to its unique characteristics and potential impact on human health. Unlike nuclear DNA, which undergoes recombination and is inherited from both parents, mtDNA reproduces asexually and is transmitted exclusively through the maternal line without recombination (Giles, Blanc, Cann, & Wallace, 1980). Despite its small size, mtDNA plays an important role in cellular energy metabolism, implicating its dysfunctions in metabolic and degenerative disorders (Lin & Beal, 2006; Lima, Li, Mottis, & Auwerx, 2022). MtDNA This distinct mode of inheritance allows mtDNA to pass unchanged from mother to offspring, making it a critical marker for genetic studies (as well as forensic identification). Although mtDNA has been implicated in a variety of aging-related and neurodegenerative disorders, including Alzheimer’s disease (Cha et al., 2012; Reddy, 2009), Parkinson’s disease (Jin et al., 2014; Winklhofer & Haass, 2010), Amyotrophic lateral sclerosis (ALS; Israelson et al., 2010; Murakami et al., 2007), and Huntington’s disease (Ayala-Peña, 2013; Oliveira & Lightowlers, 2010), a comprehensive and empirically validated model has yet to link mtDNA expression to age-related disorders (Cha, Kim, & Mook-Jung, 2015; Lin & Beal, 2006; Burt, 2023).

The singular inheritance pattern of mtDNA complicates the detection of its effects using traditional behavior genetic methods. Behavior genetic models were developed for nuclear DNA, resulting in these models being ill-equipped to separate the effects of mtDNA from nuclear DNA. When such designs are employed, mtDNA is perfectly confounded with the shared environment because siblings in these designs (full siblings, identical twins, etc.) share the exact same mtDNA. This gap highlights the need for innovative approaches that can isolate mtDNA’s influence on phenotypic traits effectively.

To address these challenges, we developed a novel covariance structure model to quantify the impact of mtDNA differences on individual differences in human traits, such as aging-related diseases (Burt, 2023; Burt et al., 2024). This model is designed to estimate mtDNA contributions by comparing phenotypic covariance among matrilineal-linked relatives to those without an unbroken chain of maternal relatives within large family datasets, while controlling for classic factors such as additive genetic variance and environmental variance. As all relatives on the matrilineal line in a family share 100% of their mtDNA and relatives on the patrilineal share 0%, we posit that the increased similarity of matrilineal relatives indexes mtDNA effects on the outcome (see Figure 1 for a visual demonstration). Statistically, the covariance structure model estimates multiple sources of influence on the variance of outcomes:

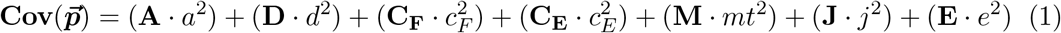

**Figure 1.**
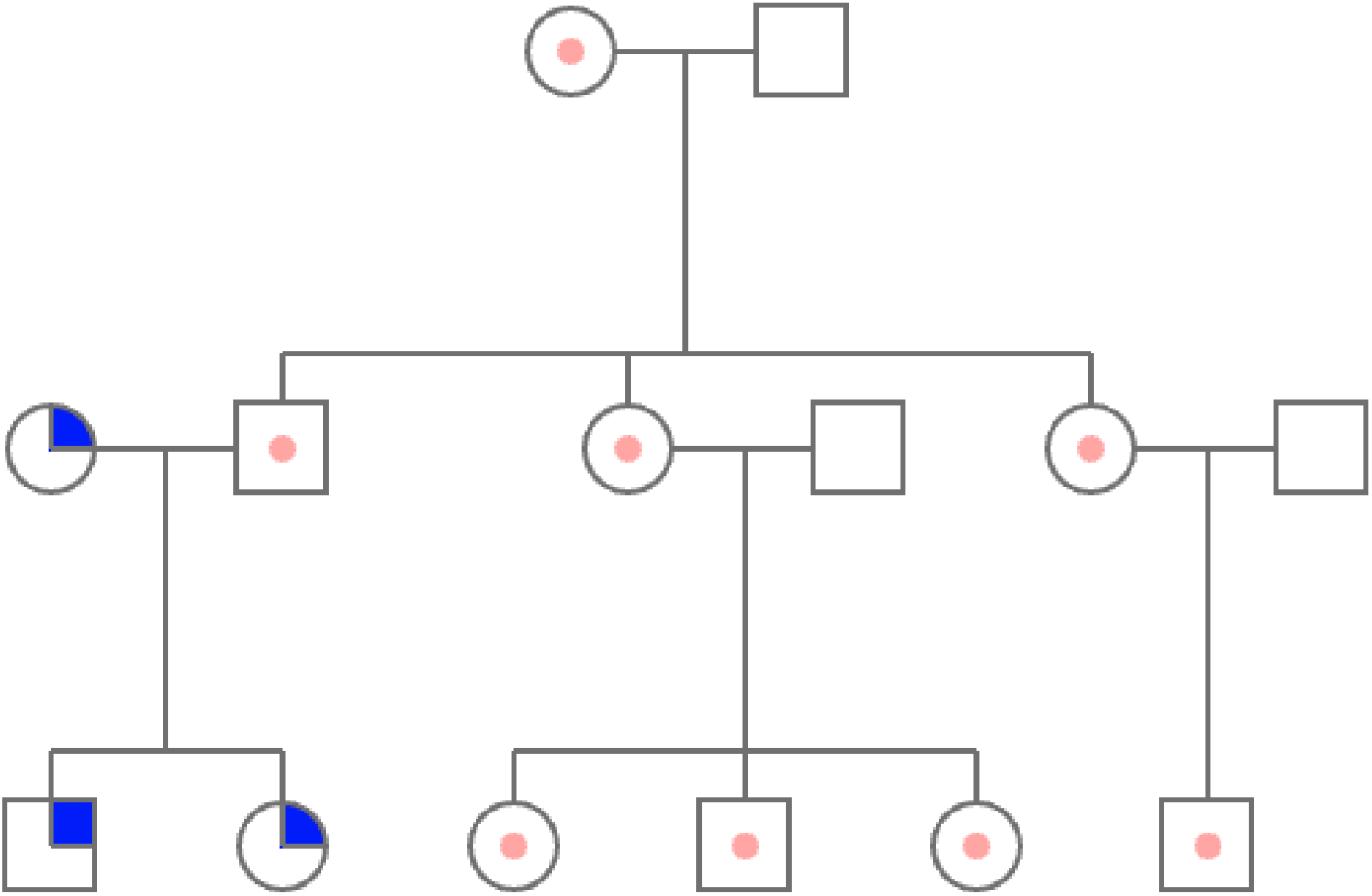
Example pedigree illustrating mtDNA inheritance. Squares denote male family members, and circles denote female members. Individuals with identical notation symbols theoretically share the same mtDNA.

In equation 1, 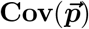 is the estimated *N × N* phenotypic outcome covariance matrix, where *N* is the number of people who have at least one non-zero relationship with any other person. The uppercase letters denote relatedness matrices, which are known and fixed in the data structure and indicate the degree of relationship between two individuals. The lower-case letters are path coefficients regressing latent variance components on the observed phenotypic variable. After being squared, these letters indicate standardized variance components that are freely estimated. **A** stands for additive nuclear DNA relatedness matrix. The *a*^2^ coefficient represents the estimated amount of variance associated with the additive genetic relatedness pattern ^1^. **D** stands for nuclear DNA epistasis relatedness matrix, calculated by element-wise squaring the additive genetic matrix. The mitochondrial relatedness matrix is abbreviated as **M**, and its corresponding variance is *mt*^2^. For kin pairs with shared maternal ancestry, it will be indicated as one, and zero otherwise. **J** stands for the interrelated regulation between nuclear and mitochondria, which is calculated by multiplying the corresponding matrix entries in **A** and **M. *C***_***N***_ indicates the matrix of the common nuclear family environment similarity. It will be coded as one if the members share the same mother and father. ***C***_***E***_ is the similarity due to the common extended family, where every member in the same pedigree will be coded as one. **E** is the unique environment matrix and random error (Burt, 2023).

The proposed model represents a pioneering approach to quantifying the impact of mtDNA on human traits. Multiple biological pathways have been found to link mtDNA differences with phenotypical differences (Ferreira & Rodriguez, 2024). The primary effects come from the transcription and translation of the 37 mtDNA encoded genes (Anderson et al., 1981). The encoded genes are translated into rRNA, tRNA and polypeptides that serve the important functions of energy production through Complex I to V and cell cycle regulations through mitochondrial-derived peptides (Moraes et al., 2002; Gruschus, Morris, & Tjandra, 2023). Another important region of mtDNA is the non-coding region and the displacement loop within the non coding region, which regulates mtDNA transcription and produces vital proteins for oxidative phosphorylation (Nicholls & Minczuk, 2014; Ferreira & Rodriguez, 2024). Both processes mentioned above contribute to the phenotypic variance that can be captured by the *mt*^2^ component of our model. One other biological property of mtDNA is that the number of copies of mtDNA within a cell differs between different individuals (Guyatt et al., 2019). There has been some preliminary research suggesting that the number of copies of mtDNA is associated with several molecular processes and human diseases such as dementia (Castellani, Longchamps, Sun, Guallar, & Arking, 2020; Chong et al., 2022). Both twin analysis and GWAS suggest that the copy number of mtDNA is regulated by nuclear DNA and influenced by environmental factors (Xing et al., 2008; Chong et al., 2022). Therefore, we expect the individual differences caused by the difference in copy number of mtDNA to be accounted for under 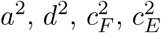 and *e*^2^.

One advantage of the current model is its ability to isolate mtDNA effects from the well-studied influences of nuclear-genetic and environmental factors. Conversely, in addition to estimating the effects of mtDNA that are not masked by other factors, the current model can also be used to estimate the effects of nuclear genetics and the environment free from any potential confounds from mtDNA effects. The current model offers the advantage of harnessing large pedigrees, allowing for the simultaneous estimation of multiple parameters. This capability enhances the robustness and accuracy of our analyses. The path tracing rules for deriving the relatedness matrices provide a solid foundation for leveraging unlimited amounts of kin pairs in pedigrees of various sizes, from small nuclear family pedigrees to pedigrees with more than 750,000 individuals (Burt et al., 2024). Previous family designs have been limited to nuclear families and extended twin designs, which has hindered researchers’ ability to estimate more variables due to a lack of degrees of freedom from additional pairwise kinship in extended pedigrees (Hunter, Garrison, Burt, & Rodgers, 2021).

Although mathematically and theoretically promising, the data used for fitting such a model faces stringent requirements. To obtain estimates of mtDNA effects, the data require family pedigrees with various unique relatedness patterns on the seven parameters. Furthermore, the family data should include a record of relevant individual outcomes. For example, since many neurodegenerative illnesses manifest at relatively late periods of life, the data must span decades for the disease to theoretically manifest in all involved generations.

### Statistical Power of MtDNA Effects Estimation

Statistical power is the probability of correctly rejecting the null hypothesis (Cohen, 1988, 1992). As the extended pedigree model is a covariance structure model, its power analysis follows the general framework of power analysis for structural equation modeling (SEM). To test the significance of a specific variance component (e.g., *mt*^2^) in a covariance-based structural equation model, the null hypothesis is that the best-fit covariance structure is the one without the specific variance component (e.g., a model without *mt*^2^). The alternative hypothesis is that the best-fit covariance structure is the one with the specific variance component (e.g., a model with *mt*^2^; Satorra & Saris, 1985). Complex covariance-based structure models are commonly fit with maximum likelihood estimation (Rao, Vogler, McGue, & Russell, 1987), which primarily employs the likelihood ratio test for hypothesis testing (Curran, Bollen, Paxton, Kirby, & Chen, 2002; Jobst, Bader, & Moshagen, 2021; MacCallum, Browne, & Sugawara, 1996; Satorra & Saris, 1985). Thus, the power to detect the estimated parameter is essentially the power of the likelihood ratio test when comparing the model under the alternative hypothesis against the model under the null hypothesis (Satorra & Saris, 1985). If the difference in log-likelihood ratio between the two models is statistically significant, we will regard that the model constructed from the alternative hypothesis fits significantly better than the model constructed from the null hypothesis.

Given the small scale of mtDNA relative to nuclear DNA (16.6k base pairs of complementary nucleotide versus 2,875,002k base pairs of complementary nucleotide; Marande & Burger, 2007), a safe assumption to be made is that the effect size of mtDNA will be substantially smaller than that of nuclear DNA for most behavioral outcomes. Previous studies have observed unusually large shared genetic influence for heights between maternally-linked cousins, which may be attributed to the impact of mtDNA (Garrison et al., 2018; Rodgers et al., 2019), but there are no previous studies directly quantifying overall mtDNA effects. Moreover, the existence of mtDNA effects is likely to vary from outcome to outcome. Hence, a power analysis should be designed and conducted to evaluate the sample size requirement under different conditions (i.e., various family structures; different sets of estimated parameters; different effect sizes of the mtDNA effects).

Since the current covariance structure model is an extension of a standard twin-family ACE model, factors that impact the power of the ACE model could serve as a heuristic starting point for the current model. Previous studies have thoroughly discussed how sample size (Martin, Eaves, Kearsey, & Davies, 1978; M. Neale & Cardon, 2003; Verhulst, 2017; Visscher, 2004), variance components (Verhulst, 2017), the type-one error rate (Sham, Purcell, Cherny, Neale, & Neale, 2020), and the ratio of twin types (Visscher, 2004; Visscher, Gordon, & Neale, 2008; Lyu & Garrison, 2023) impact the power of parameter estimation in the ACE model. In the case of mtDNA effects, it is reasonable to hypothesize that some properties of the family structure will impact the power estimation, given the fact that the power of the ACE model is also impacted by the ratio of MZ and DZ twins and the distribution of variance contribution (Martin et al., 1978; Visscher, 2004). Visscher (2004) found in every variance combination of the ACE model, there exists an optimal ratio of MZ over DZ for a given sample size to reach a specific power. For example, when the population variance components are distributed as *a*^2^ = .6, *c*^2^ = .1, to achieve a power of .95, a total of 235 pairs of twins is required, of which 70% should be MZ twins. For the mtDNA effect, power can be potentially impacted by the proportion of matrilineal relatives (relatives who share the identical mtDNA in a family) and patrilineal relatives (relatives who do not share the same mtDNA in a family). Just as there exists an optimal ratio for MZ and DZ twins in ACE model estimation, an optimal ratio between the matrilineal and patrilineal relatives should exist for mtDNA effect estimation. However, in reality, researchers have very limited control of the gender distribution in pedigree datasets. In all our simulations, we fixed the gender ratio at biological male : biological females at 1:1.

Besides the ratio, families with different structures inherently have different relatedness matrices. Including additional unique relatedness matrices in the model will provide more degrees of freedom for estimation. A previous study on covariance-based structural equation modeling (SEM) indicated that having more degrees of freedom can enhance the power of the estimation (Maccallum, 1996). However, for a fixed amount of individuals, larger families will lead to a decrease in the number of involved families, which results in a shrink in the amount of relatedness matrices. Many properties of a pedigree, such as the number of sibships in each family and the number of generations involved, will impact the scale of each relatedness matrix that can be derived (Chen & Abecasis, 2006; Hunter et al., 2024). Conducting systematic research on how pedigree structures impact mtDNA effect estimation power will provide useful guidelines for future users of the current model. It will contribute to the field’s understanding of the incorporation of extended kinships beyond siblings and nuclear families, as such designs have demonstrated effectiveness in reducing biases inherent in twin/sibling studies and in offering fresh perspectives (Keller, Medland, & Duncan, 2010). In family pedigrees, the number of generations involved (the depth of a pedigree) and the average number of offspring per couple (the width of a pedigree) prominently determine the overall shape of the pedigree and the total number of family members. In addition to the factors mentioned above, the proportion of fertile spouses among all the spouses and the relative ratio of biological sexes also influence the pedigree’s shape. We define the factors as the mating rate and newborn sex ratio of the pedigree, respectively. Despite the importance of mating rate and newborn sex ratio, we will mainly operationalize the pedigree shape by manipulating the number of generations and average number of offspring per couple in our paper.

### Model Assumptions and Estimation Bias

Like other family structures, unbiased estimation of mtDNA effects relies on ensuring certain assumptions to avoid model misspecification. For instance, in classical twin studies, the model’s robustness depends on assumptions such as random mating, which validates the 0.5 genetic relatedness in dizygotic twins, and the equal environment assumption, indicating that both MZ twins and DZ twins share family environments to the same extent (Felson, 2014; M. Neale & Cardon, 2003). However, applying the extended pedigree model introduces several concerns:

1. Firstly, the estimation can be impacted by unobserved kinship in large datasets, as estimating mtDNA effects requires extended pedigrees. For example, if two nuclear families are connected through the same mother, but one family lacks documentation of this maternal relationship in the dataset, the mtDNA relatedness matrix will inaccurately contain zeros where there should be ones. Such missing links can significantly affect the accuracy of mtDNA effect estimation. No family dataset is collected perfectly, and all of them have unidentified or incorrectly identified kinship links to some extent, depending on the data collection practice. Hence, testing the robustness of the estimation given an imperfect kinship record ensures the efficacy of the model as a pivoting method on detecting mtDNA effects.
2. Secondly, the current model assumes no mutation of mtDNA across generations. Like nuclear DNA, mtDNA mutates randomly during replication and the stability of polynucleotide chains decreases under certain external influences(DiMauro & Schon, 2001). Given that mtDNA comprises only about 17k base pairs and 37 encoding genes, any mutation could lead to a misspecification of mtDNA relationships among individuals sharing the same matrilineal line in a pedigree. However, in most datasets where this model will potentially be applied, actual mtDNA sequencing is unlikely to have been performed. Given a mutation rate of 2.7Ö10^*-*7^ per site per generation, we estimate that the expected number of mutations per generation is around 4.59Ö10^*-*3^ in germline cells. In other words, the mutation rate suggests that one mutation is expected to happen every 218 parent-offspring transmissions. Although the absolute rate of mutation rate is low, if the undetected mutation exists between the early generations of the pedigree, the mtDNA relatedness matrix will be misspecified for a lot of decedents after the mutation. Hence, understanding how the estimates of *mt*^2^ will be impacted by this limitation is important for future applications.
3. Thirdly, unlike the nearly universal additive genetic effects observed across human traits, mtDNA’s influence may be minimal for certain traits. Therefore, assessing the false positive rate of the estimation is crucial to ensure that variances from other components are not mistakenly attributed to mtDNA effects, when there are no mtDNA effects for the specific traits.

## Methods

In this study, we employed Monte Carlo simulations to investigate two critical statistical properties of the extended pedigree model. First, we examined how statistical power for mitochondrial DNA (mtDNA) estimation varies across different sample sizes and explored the influence of diverse pedigree structures on power. Second, we evaluated the effects of specific model assumption violations on estimation bias, quantifying both the direction and magnitude of the bias under each scenario. All simulation R code for this paper is available at https://github.com/Xuanyu-Lyu/mtDNA, along with the accompanying R package (BGmisc; Garrison, Hunter, Lyu, Trattner, & Burt, 2024).

### Power Analysis

In the current analysis, we operationalize the power of mtDNA estimation as the power to detect *mt*^2^ and *j*^2^ simultaneously, as both variances are vital components of the mtDNA effects. In the current study, our null (*H*_0_) and alternative hypothesis (*H*_1_) are:

*H*_0_: The mtDNA (*mt*^2^) and its interaction with nuclear DNA (*j*^2^) do not have a non-zero effect on a specific phenotype.

*H*_1_: The mtDNA (*mt*^2^) and its interaction with nuclear DNA (*j*^2^) have a non-zero effect on a specific phenotype.

To evaluate the significance of the likelihood ratio test, a unit test statistic (*T* ) is computed by comparing the fits of two models: the null hypothesis model and the alternative model. The test statistic is calculated as the difference in their respective negative twice- log-likelihood values, normalized by the total sample size (*N* ) used in the simulation:

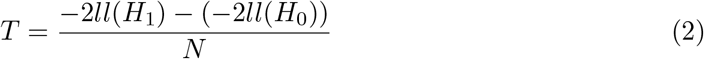

Here, *ll*(*H*_0_) and *ll*(*H*_1_) represent the negative twice-log-likelihood values derived from fitting the null hypothesis model and the alternative model, respectively, while *N* denotes the total number of individuals included in the simulation.

The test statistic *T* follows a non-central chi-square distribution:

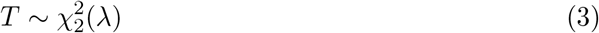

where 2 is the degrees of freedom of the distribution and the non-centrality parameter (*λ*) is derived from

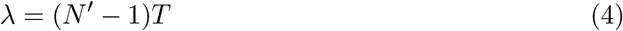

where *N* ^*′*^ is the target sample size that we want to derive power from.

Then, the statistical power of the likelihood ratio test given any sample sizes can be derived by comparing the central *χ*^2^ distribution from the null hypothesis and the non-central *χ*^2^ distribution from the alternative hypothesis:

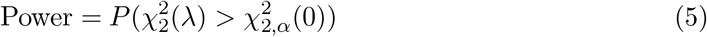

where 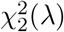 is a random variable following a non-central chi-square distribution with 2 degrees of freedom and non-centrality parameter *λ*, and 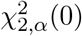 is the critical value from the central chi-square distribution at significance level *α*. In the current study, *α* = .05.

One convenient property of power analysis for the likelihood ratio test is that only one robust unit test statistic *T* , which is the contribution to *λ* from one individual, is required. Afterward, power for any sample sizes under the same condition can be easily derived numerically using equations (4) and (5). The essential unit test statistic (*T* ) for different pedigree structures is obtained by averaging *T* observed from 100 simulation sets, each including 10,000 individuals. The association between sample sizes and power for various conditions can be easily derived when an accurate non-centrality chi-square parameter is obtained through simulation (Satorra & Saris, 1985; Verhulst, 2017).

For family pedigrees, various factors such as the number of generations involved, the average number of offspring per mate, sex ratios of newborns, mating rates within a generation, inbreeding, remarriage rates, and adoption rates significantly influence the structure and size of the family pedigree. As noted, the depth (number of involved generations) and the width (average offspring per couple) of a pedigree are also critical in shaping the overall family structure and determining the total family size. To investigate the impact of these two factors, we set up five conditions of parameters that influence family structure (Table 1). Among these conditions, pedigrees 1, 2, and 3 are designed for evaluating the impact of the number of generations, and pedigrees 1, 4, and 5 are designed for evaluating the impact of offspring per mate. Due to the relatively low prevalence of factors like inbreeding and twins, we did not perform power analysis under variation of other pedigree structure parameters. We did not analyze the impact of other factors including sex ratios and mating rates pedigrees on power, since their impact is rather limited in a real-life setting, and researchers have very little control over these factors. We fixed sex ratios of newborns at 1:1 and the mating rates at .7. We also assumed these low-prevalence scenarios like consanguinity and twins did not exist in our pedigrees to reduce design complexity and increase computational efficiency.

**Table 1.**
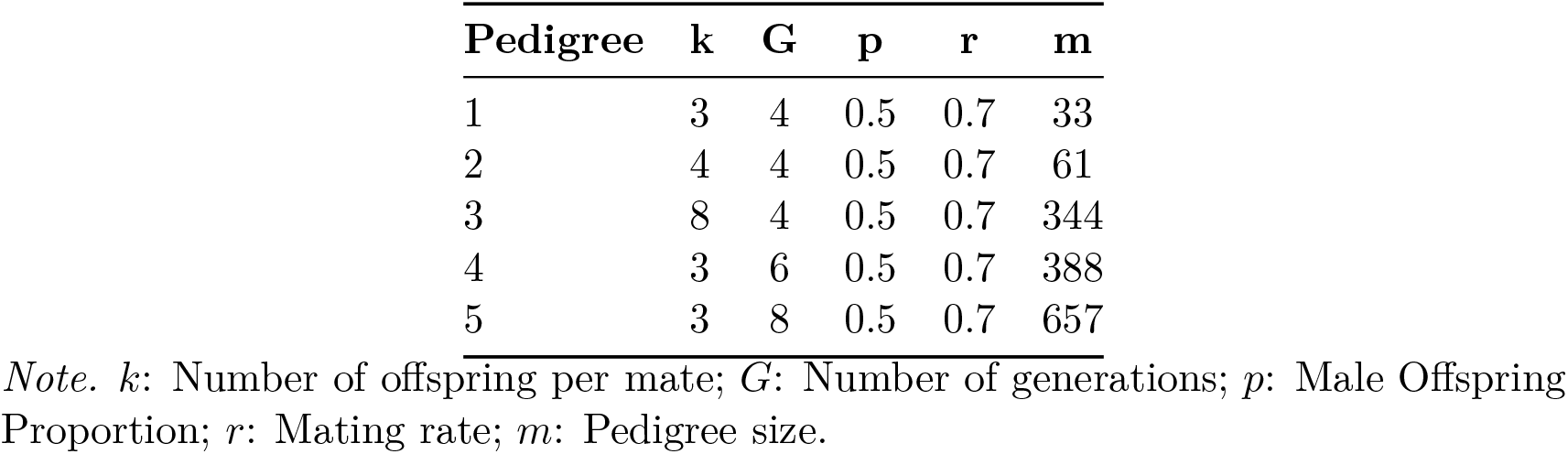
Pedigree Structures Used in Simulation.

Family size is another important factor in family structure. In the current study, we did not set family size as a self-varied parameter. The family size of one specific condition is a weighted exponential function of the number of generations, average offspring, and mating rate under the current assumptions of family structure. In equation 6, for any *g ∈* ℤ

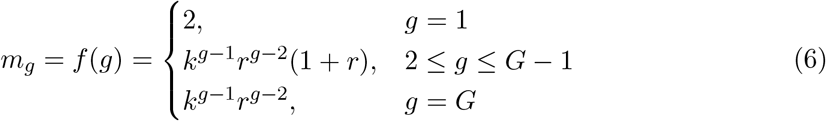

where *m*_*g*_ is the number of individuals in the *g*^*th*^ generation, *k* is the number of offspring per mate, *r* is the mating rate, and *G* is the total number of generations. The number of individuals *m* in a pedigree with *G* generations can be calculated with 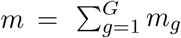.

We developed a simulation function that generates random pedigree structures based on user-defined parameters, ensuring variability in the simulated data to more closely resemble real-world scenarios (Garrison et al., 2024). In this model, the number of offspring for a given pairing is determined by a Poisson distribution, where the mean is set to the user-specified average number of offspring. This approach allows for variation within each generation, although the total number of generations remains constant, as dictated by the user’s parameters. The exact number of generations in a simulated pedigree is fixed at the value of the user-specified parameter for the convenience of deriving the non-centrality parameter, as ideally, the size of the pedigree should remain constant. All simulated pedigrees will start with an identical 1st generation that only has one mated couple. The pedigree will end with a generation in which no individual has mated. For instance, figure 2 presents two example pedigree plots from the same parameter setting (*G* = 3, *k* = 4, *r* = .7) of the simulation function. Both pedigrees have four generations, but the number of offspring across mates are different, while the size of the pedigree remains almost identical.

**Figure 2.**
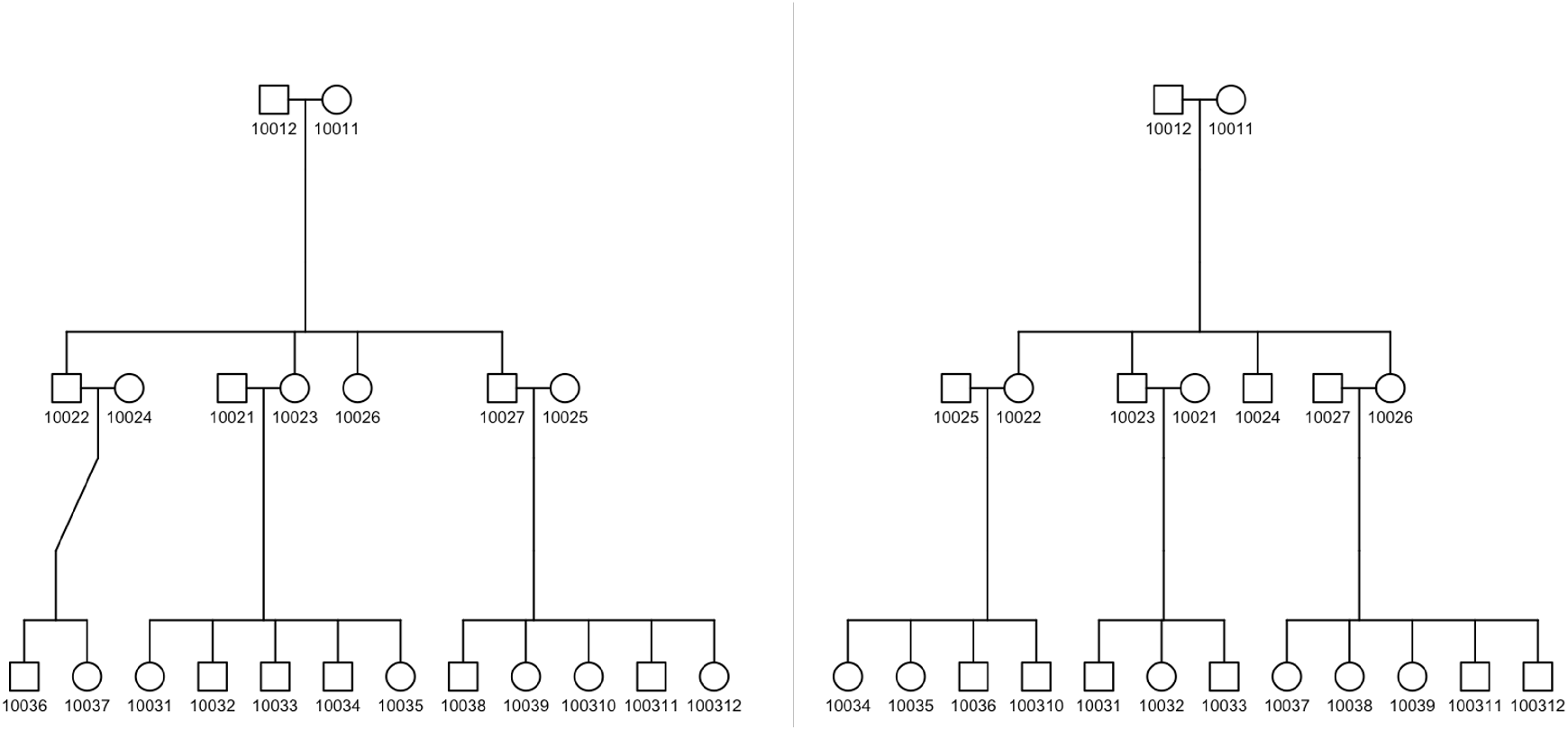
Two pedigrees generated with the same parameters: Number of generations (*G*) is 3, number of offspring per mate (*k*) is 4, and male offspring proportion (*p*) is 0.7, using the *simulatePedigree()* function from the BGmisc R package.

In our simulations, we introduce four distinct levels of *mt*^2^ estimates to explore the sample sizes necessary to detect various levels of mtDNA effects. The specified variance levels explained by mtDNA are set at 0.5%, 1%, 5%, and 10%. We keep other variance components constant across all conditions, as they do not constitute the primary focus of this study. Specifically, we define *a*^2^ = .40, *d*^2^ = 0, 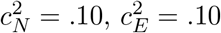, and *j*^2^ = .01. Indicated by previous research on the ACE model (Visscher, 2004; Verhulst, 2017), alterations in one variance component can influence the estimations of others. Consequently, the power to detect *mt*^2^ and *j*^2^ variations is likely influenced by the levels of *a*^2^ and 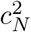.

### Estimation Bias When Violating Assumptions

In the current study, we analyzed the estimation bias that arises when various model assumptions are not met. We used the BGmisc package (Garrison et al., 2024) to simulate data that did not meet the assumptions and then fit the model assuming that the assumptions were met. Our goal was to determine if the estimates of mtDNA effects remained robust even when the data did not meet the model assumptions, or when we were uncertain if the data met the assumptions. All simulations conducted to evaluate estimation bias use a variance combination of *a*^2^ = 0.40, *d*^2^ = 0, *mt*^2^ = .05, 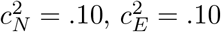, and *j*^2^ = .01.

The first potential source of estimation bias we examined was the assumption that the pedigree measure in the data is intact, which is unlikely to be true in most real-world pedigree datasets. These deviations can be considered as random noise in the pedigree structure. Incomplete sampling of a pedigree can lead to the separation of relatives that are originally from the same pedigree. When estimating the effects of mitochondrial DNA (mtDNA), it’s important to note that some maternal relatives may not be identified in the kinship records. This can happen if there is a missing link in the true pedigree that connects a female individual to her relatives, such as a mate or parent-offspring relationship. This can result in relative pairs who have all genetic and environmental relatedness matrices being misspecified as not related, potentially leading to biased estimates of mtDNA effects. To examine bias, we simulated 500 sets of 124 pedigrees (*n* = 161; *k* = 3; *G* = 5; *r* = 1, and total sample size of each simulation *N* = 19964). Phenotypic data were generated based on relatedness matrices derived from the true complete pedigree. Next, we randomly selected a female relative in the second generation and dropped her kinship with her mother and father, resulting in misspecified relatedness matrices. We repeated the process for all 10,000 simulated pedigrees. The model was fitted using correct and incorrect relatedness matrices, resulting in two sets of estimates of mtDNA effects. By comparing the distribution of 500 estimates in both the correct model and the misspecified model, we can evaluate the bias of the estimates when there are unobserved maternal links in the pedigree.

The second source of potential bias we evaluated was the unmeasured presence of mtDNA mutations. If a female relative in a pedigree has a mutation in her mtDNA when forming her gametes, her descendants will have a mtDNA relatedness coefficient of 0 with all other individuals in the pedigree who do not share the same mtDNA mutated female ancestor. In other words, the mutated individual and all his/her maternal progeny will have a different haplotype of mtDNA from the individual’s maternal ancestors. This is consistent with how we designate shared and non-shared mtDNA haplogroups in the mtDNA relatedness matrix initially. In our model, we assume that all maternal relatives in a pedigree share the same mtDNA. However, this assumption is incorrect when a mutation occurs and will lead to a misspecified mtDNA relatedness matrix. The missing mutation can cause biased estimation of mtDNA effects. We thus simulated 500 sets of 124 pedigrees (*n* = 161; *k* = 3; *G* = 5; *r* = 1, and total sample size of each simulation *N* = 19964). Then, we randomly selected a female relative in the 2nd generation and dropped the female’s kinship with her parents. We generated the mtDNA relatedness matrix with the mutated pedigree, but all other relatedness matrices are generated from the intact pedigree. We examine the bias from not incorporating mutations by comparing the distribution of 500 estimates from the correct model (with mutation) and the model of which mtDNA relatedness matrix is misspecified (without mutation).

Third, we investigate the impact of applying the full covariance structure model with seven parameters to phenotypes that are not influenced by mtDNA (i.e., *mt*^2^ = 0, *j*^2^ = 0). By fitting the full model with the simulated data, we can evaluate the proportion of the 500 estimates that have values exceeding a certain numerical value as a metric of the false positive rate of the *mt*^2^ estimate. We simulated 200 sets of 124 pedigrees (*n* = 161; *k* = 3; *G* = 5; *r* = 1, and total sample size of each simulation *N* = 19964) and generated data using a covariance matrix without the contribution of mtDNA effects.

## Results

### Power

In Table 2, we present the sample size required to reach a specific power level under various effect sizes of *mt*^2^. To compare the effects of pedigree structure, we designated pedigree 4 as a large pedigree and pedigree 1 as a small pedigree. If not specified, we used a variance combination of *a*^2^ = 0.40, *d*^2^ = 0, *mt*^2^ = .05, 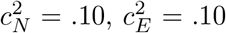, and *j*^2^ = .01 for all other parameters. In general, 30,000 individuals will grant an adequate power when 1% or more of the phenotypic variance is explained by mtDNA. However, large sample sizes are required for traits under minor mtDNA effects, especially when using smaller pedigrees.

**Table 2.** Power Analysis for Different Pedigree Sizes and Variance Levels.

| Pedigree size | Variance level | Power |  |  |  |
| --- | --- | --- | --- | --- | --- |
|  |  | 0.4 | 0.6 | 0.8 | 1.0 |
| Large pedigree (m = 388) | $mt^2 = .1$ | 576 | 933 | 1447 | 5478 |
| | $mt^2 = .05$ | 1861 | 3017 | 4678 | 17708 |
| | $mt^2 = .01$ | 11671 | 18919 | 29340 | 111073 |
| | $mt^2 = .005$ | 14630 | 23717 | 36780 | 139240 |
| Small pedigree (m = 33) | $mt^2 = .1$ | 607 | 984 | 1526 | 5775 |
| | $mt^2 = .05$ | 2038 | 3304 | 5124 | 19396 |
| | $mt^2 = .01$ | 12312 | 19959 | 30953 | 117179 |
| | $mt^2 = .005$ | 13995 | 22687 | 35183 | 133193 |

*Note.* The table presents the sample size (number of individuals) required for specific power levels to detect genetic variance at different $mt^2$ levels in large and small pedigrees.

#### Pedigree-Number of Generations

In Figure 1, three pedigree structures with varying numbers of generations (*G*) are compared in terms of power to detect mtDNA effects. In these pedigrees, each mate has three offspring (*k* = 3). The findings from Figure 3 demonstrated that pedigrees with a larger number of generations have increased power. When examining pedigrees with an eight-generation depth (*G* = 8), a power of .8 is attained with 4,011 individuals, utilizing a variance combination with medium mtDNA effects (*mt*^2^ = .05, *j*^2^ = .01). On average, across all simulated variance component combinations, using four-generation pedigrees requires 27.76% larger sample sizes compared to five-generation pedigrees in order to achieve a power of .8. Conversely, the smallest pedigrees (*G* = 4) exhibit the lowest power, necessitating a sample size of 5,124 individuals to achieve a power of .8.

**Figure 3.**
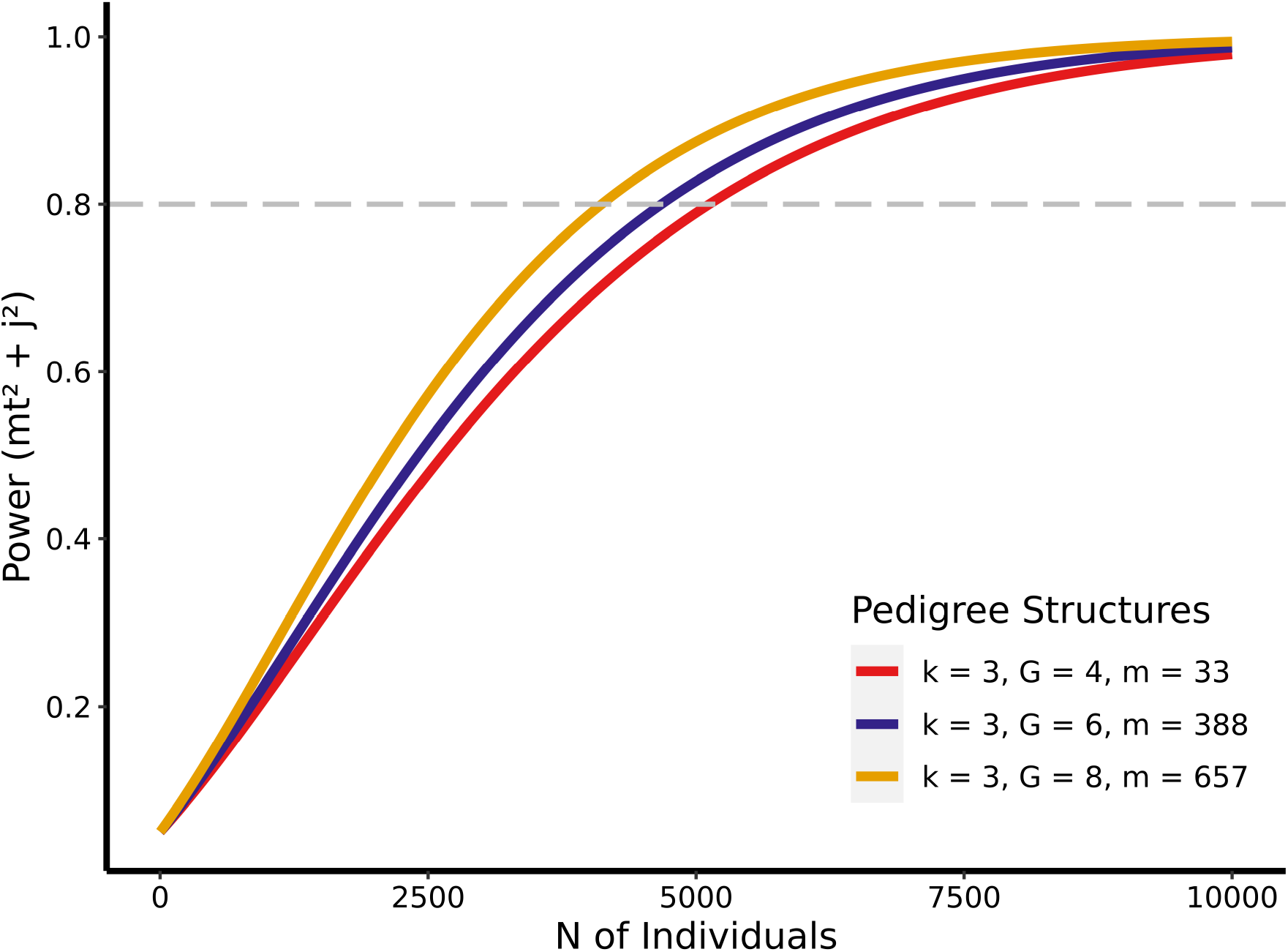
Estimation power for *Mt*^2^ and *J* ^2^ as a function of total sample sizes and number of generations in simulated pedigrees. *Note: k*: Number of offspring per mate; *G*: Number of generations; *m*: Pedigree size. Effect sizes of variance components: *a*^2^ = 0.40, *d*^2^ = 0, *mt*^2^ = .05, 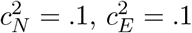, and *j*^2^ = .01.

#### Pedigree-Offspring per mate

We find that the average number of offspring per mate (*k*) has a lower influence on the estimation of mtDNA effects, in comparison with the impact of number of generations (*G*). Our findings reveal that a higher number of offspring per couple leads to slightly decreased power to detect mtDNA effects, as demonstrated in Figure 4. In particular, for medium mtDNA effects (*mt*^2^ = .05, *j*^2^ = .01), a sample size of 5755 individuals from pedigrees with an average of eight offspring per mate (*k* = 8) is necessary to achieve a power of .8 when using a four-generation deep pedigree. However, when each mate has three offspring on average (*k* = 3), only 5124 individuals are needed to reach a power of .8, resulting in greater power than with a larger nuclear family.

**Figure 4.**
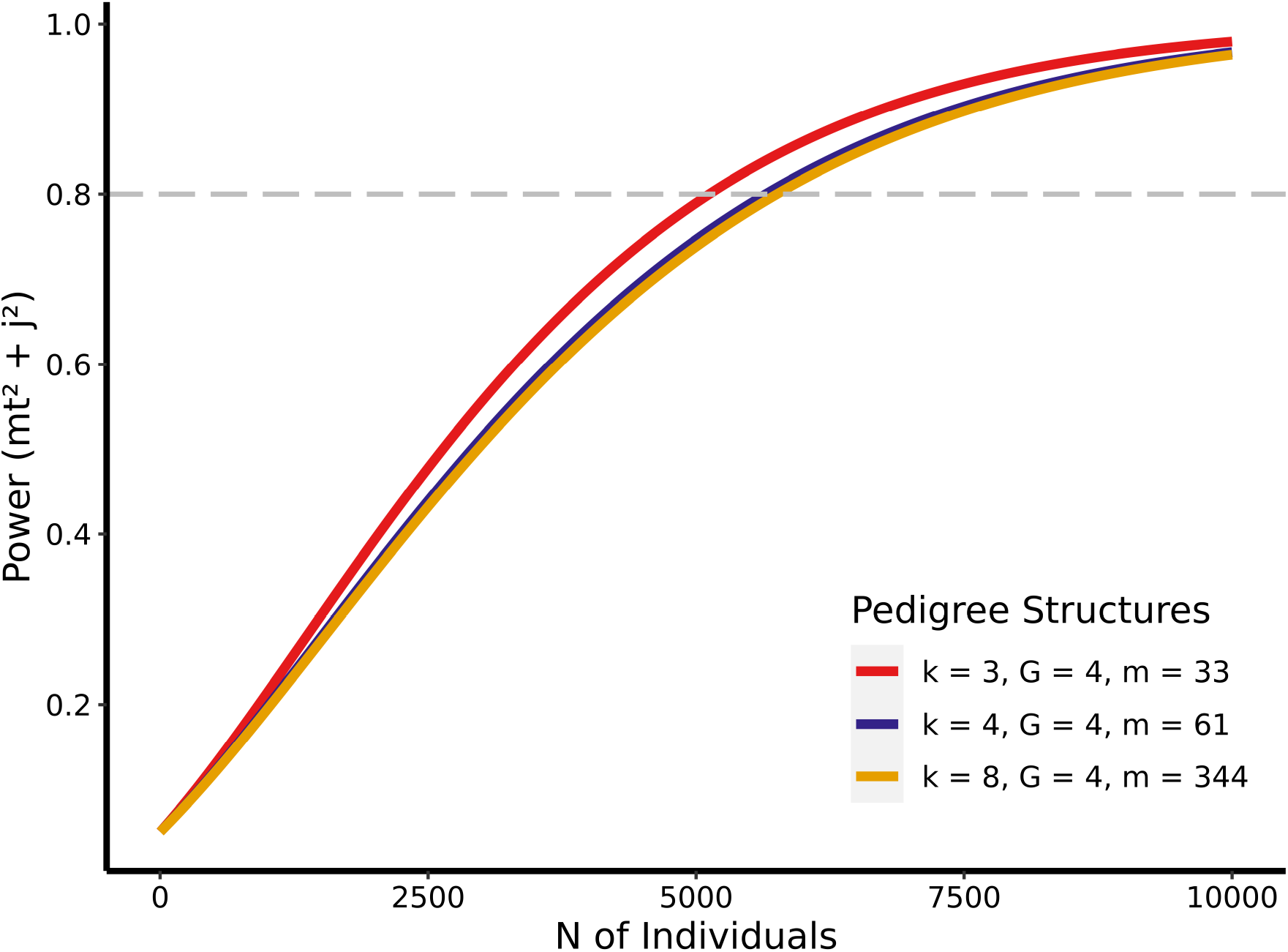
Estimation power for *Mt*^2^ and *J* ^2^ as a function of total sample sizes and number of offspring per mate in simulated pedigrees. *Note: k*: Number of offspring per mate; *G*: Number of generations; *m*: Pedigree size. Effect sizes of variance components: *a*^2^ = 0.40, *d*^2^ = 0, *mt*^2^ = .05, 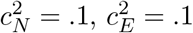, and *j*^2^ = .01.

### Model Misspecification

#### Mutation

In our simulation, we introduced a single nucleotide mutation in the mtDNA at the time of gamete formation for a single female individual in the second generation. When this mutation is not correctly specified, it leads to a cascade of misspecifications in her offspring. Across randomly simulated pedigrees, we found that *mt*^2^ is underestimated by 38.86%. This is demonstrated in Figure 5, where the mean *mt*^2^ estiates from 500 simulations decreased from .0466 to .0285 when the simulated population level was .05. Conversely, *j*^2^ parameters are drastically overestimated, deviating from .0175 to .0563.

**Figure 5.**
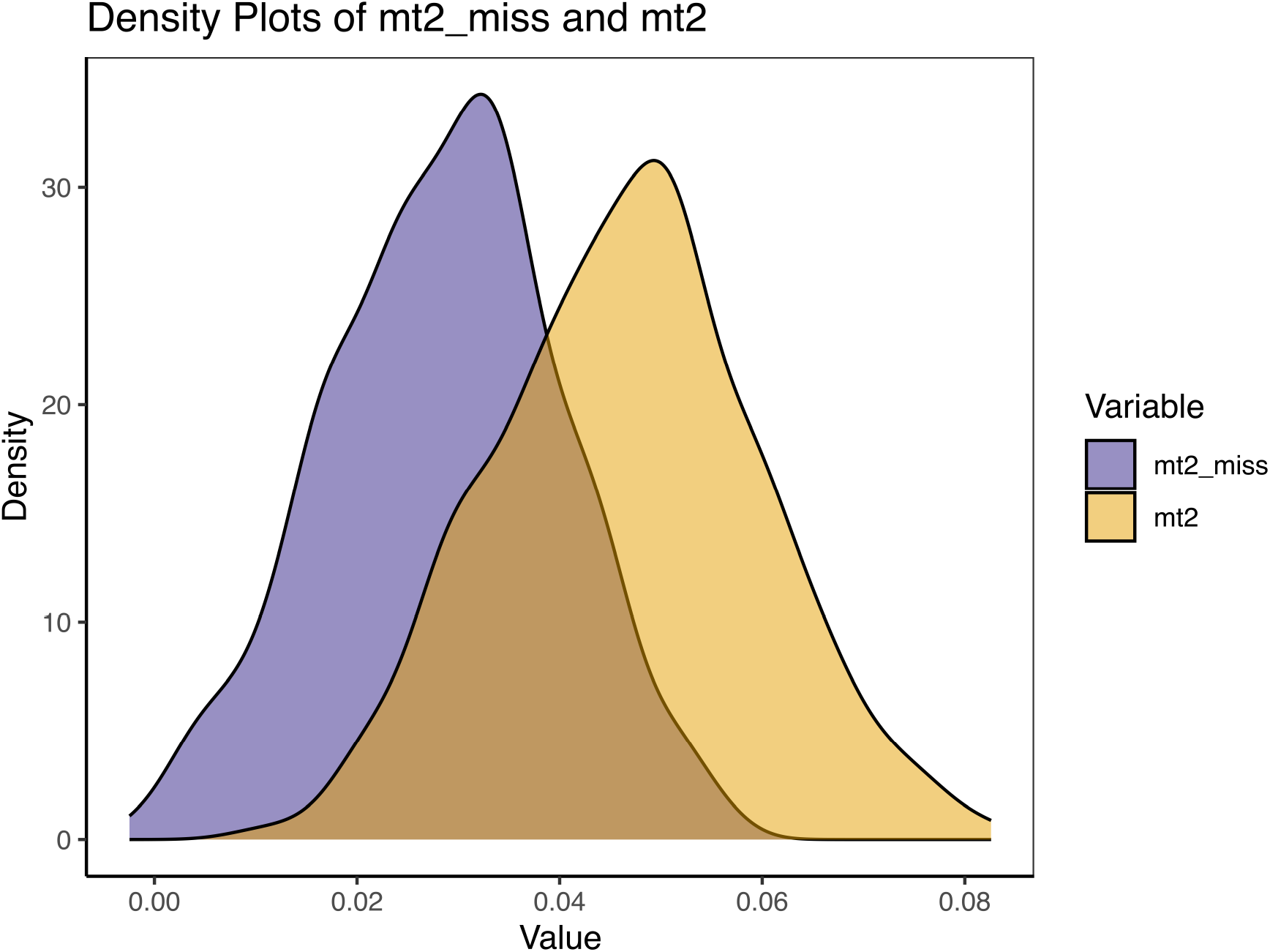
Density plot of 500 *mt*^2^ estimates. The purple shade is the biased estimation: fitting the model with relatedness matrices which includes one undetected mtDNA mutation in one second-generation female member from the pedigree. The yellow shade is the unbiased estimation: fitting the model with correct relatedness matrices.

#### Missing Links

We find that *mt*^2^ is underestimated by 38.65% if we do not correctly specify the parent-offspring relationship for a female offspring from the second generation. Demonstrated in figure 6, the mean *mt*^2^ estimates from the 500 simulations reduced from .0472 to .0289, when the simulated population level is .05. Similar to the unidentified mutation case, *j*^2^ parameters are overestimated, increasing from .0186 to .0606.

**Figure 6.**
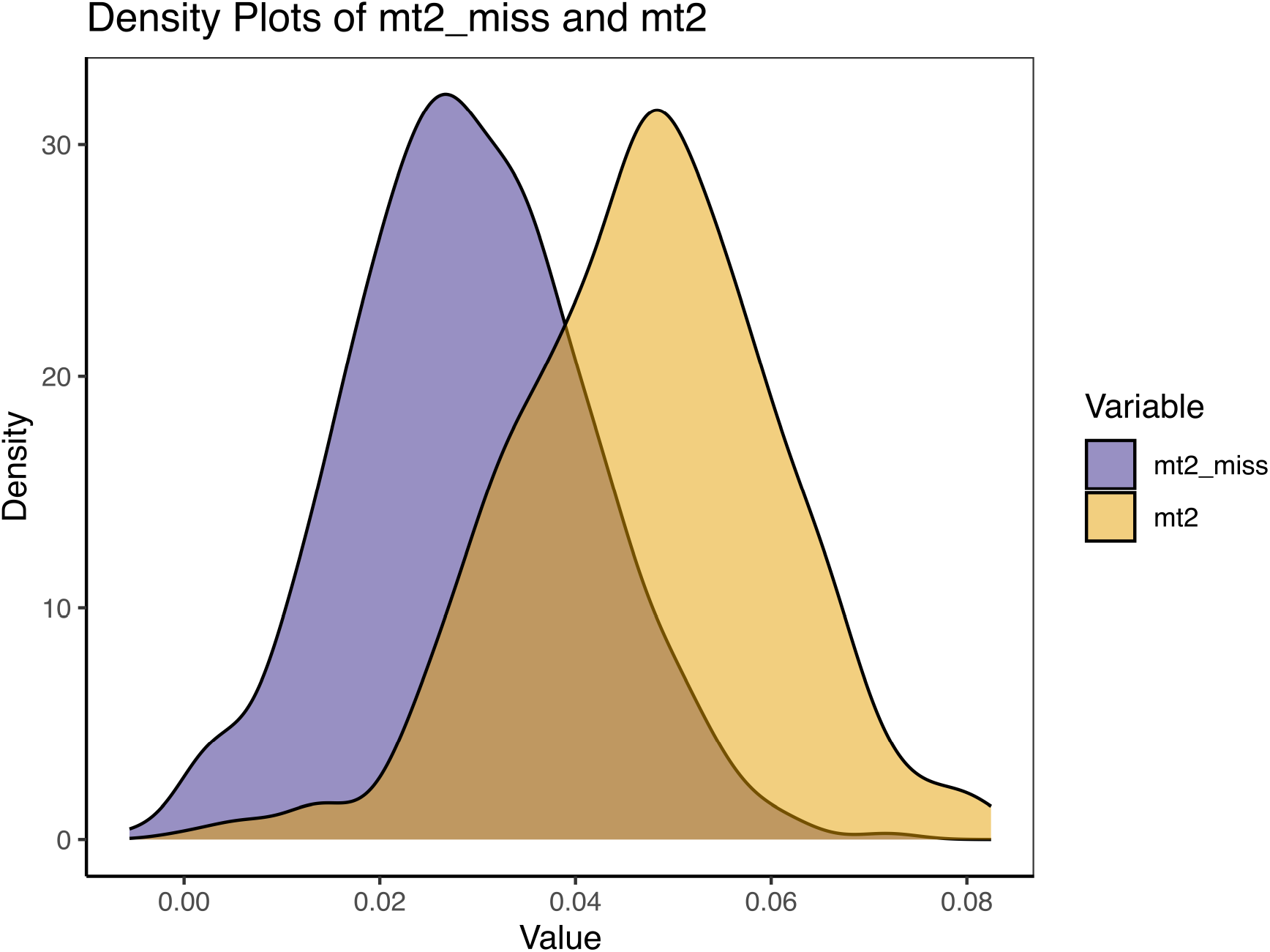
Density plot of 500 *mt*^2^ estimates contrasting biased and unbiased estimations. The biased estimation (purple shade) involves fitting the model with relatedness matrices that include an undetected mtDNA mutation in a second-generation female member of the pedigree. The unbiased estimation (yellow shade) involves fitting the model with correct relatedness matrices.

**Figure 7.**
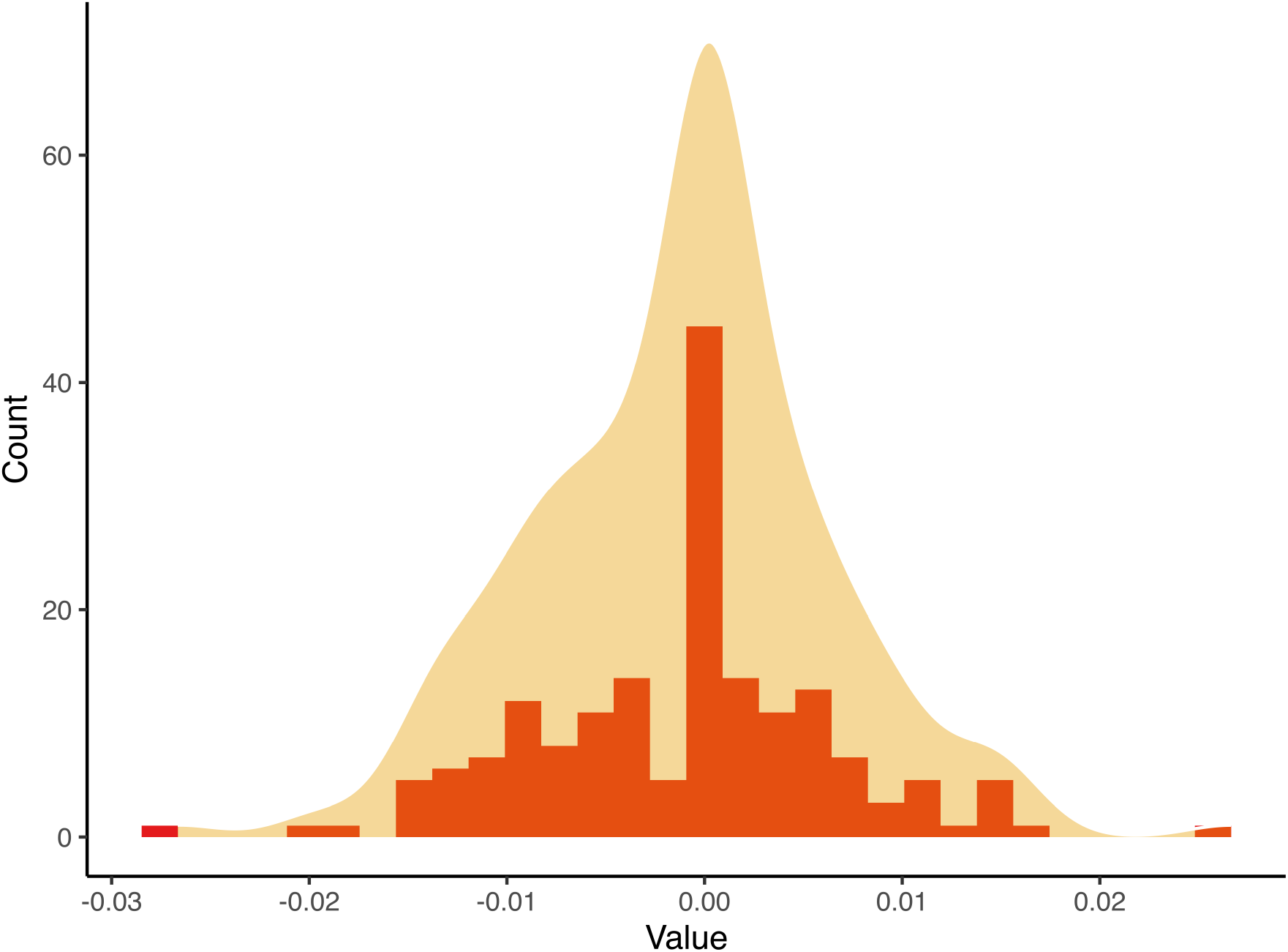
Histogram and density plot showing the distribution of *mt*^2^ estimates across 500 simulations under the null hypothesis, centered around the expected value of zero.

#### False Positives Under the Null Hypothesis

To evaluate our extended pedigree model’s performance under the null hypothesis—where no mtDNA effects (*mt*^2^ and *j*^2^) are expected — we simulated data under the null model (*mt*^2^ = 0; *j*^2^ = 0) and fit that data to the full mtDNA model. As shown in figure 5, the estimates for *mt*^2^ and *j*^2^ were unbiased and did not result in excessive false positives. Across the 200 simulated models, *mt*^2^ have a mean of -.001 and a standard deviation of .0006. *j*^2^ have a mean of .005 and a standard error of .0019. Further, 80.80% of the *mt*^2^ estimates fell within a 0.01 deviation from 0 (the true population parameter), and all were within a 0.03 deviation, while 44.07% of *j*^2^ estimates fell within the same 0.01 range. For any traits with a true *mt*^2^ estimate greater than .03, the p value for the test will be 0 when the sample size is not less than 20,000.

Overall, the results indicate that our mtDNA estimates are reasonably robust to assumption violations. In cases of extreme misspecification, the inability to detect missing maternal linkage and failure to account for mtDNA mutation led to a slightly underestimated variance of mtDNA at the population level. Furthermore, when fitting the model with data simulated under the null model, a minor false positive estimate in *mt*^2^ estimates was introduced.

## Discussion

The current study investigated the statistical power and estimation bias when using an extended pedigree model to estimate mitochondrial DNA (mtDNA) effects. Our findings suggest that a sample size of approximately 5,000 is generally adequate to detect an mtDNA effect size of 5%. We found that deeper pedigrees enhanced the model’s power to detect mtDNA effects, whereas wider pedigrees tended to diminish it. Additionally, our examination of estimation bias underscores the robustness of the extended pedigree model in estimating mtDNA effects. Specifically, failure to account for missing relatedness in the pedigree and possible mtDNA mutation both lead to slight underestimates of mtDNA variance. Overall, the false positive rate is modest for mtDNA estimation from the extended pedigree model. Collectively, these results support the extended pedigree model’s efficacy and validity for investigating the influence of mtDNA on human traits.

The power to detect mtDNA increased with larger effect sizes, as demonstrated by our simulations with mtDNA effect sizes ranging from large (*mt*^2^ = .10) to small (*mt*^2^ = .005). This trend aligns with previous power analyses for additive genetic effect estimation and common environmental effects estimated through classical twin and other family designs (Posthuma & Boomsma, 2000; Sham et al., 2020; Verhulst, 2017; Visscher, 2004; Visscher et al., 2008). Moreover, in the broader context of structural equation modeling (SEM), effect size is consistently identified as one of the most critical factors affecting power (Curran et al., 2002; MacCallum et al., 1996; Wang & Rhemtulla, 2021). Besides an increase in the effect size of targeted effects itself, reduction in error variance in general leads to higher power for the estimation of the target variance component, paralleling from ACE or ADE models which show that greater variance attributed to common environmental effects (*c*^2^) improves power for *a*^2^ estimates (Posthuma & Boomsma, 2000; Verhulst, 2017).

Our study defined pedigree width by the number of offspring per mate (*k*) and depth by the total number of generations (*G*), which are the two main factors influencing the shape of family trees. We found that a larger *G* leads to a greater power contribution per individual, but a larger *k* decreased it. In other words, data with more far-reached ancestries will grant researchers advantages in detecting mtDNA effects while a contemporary and more fertile population may be less ideal for such analysis, given the same amount of individuals in the data. The advantage of deeper pedigrees lies in their ability to more clearly distinguish between the effects of nuclear and mitochondrial effects. Conversely, wider pedigrees often yield redundant information to parse mtDNA effects from nuclear DNA. To understand the separation of *a*^2^ from *mt*^2^, we need to recognize that all members within the same generation either share a 0.5 additive genetic relatedness as siblings or a 0.125 as cousins. In contrast, between-generation kinship varies more in additive genetic relatedness. Therefore, “wide” pedigrees have more kinship links with 0.5 and 0.125 expected additive genetic relatedness, while “deep” pedigrees have links with expected values less than 0.125. At the same time, both the “wide” and “deep” pedigrees have roughly the same proportion of kinship that share the same mtDNA. Consequently, the effects of mtDNA will be easier to manifest in the “deep” pedigree because the influence on similarity between a pair of members, stemming from mtDNA, can be more easily separated from the similarity originating from nuclear DNA. This is due to the smaller average expected nuclear genetic relatedness values in the “deep” pedigrees. In other words, collecting data with more embedded generations or expanding existing datasets to include information from earlier and later generations can be a viable approach to increasing power.

The current model presupposes that kinship records are entirely accurate and that mtDNA mutations are absent. Yet, these conditions are rarely met in practice. Our bias analysis indicated that mtDNA effects are underestimated when relatedness matrices are misspecified. Given that the extended pedigree model’s aim is to pioneer the evidence of mtDNA effects on specific traits, underestimating these effects is preferable to committing a type-I error. Consequently, researchers can be confident when they uncover significant mtDNA variance, even in samples that have imperfect kinship records. Moreover, our scenarios of missing kinship and ignored mutations involve extreme misspecifications of relatedness matrices, scenarios unlikely in real-world settings. For instance, our analysis of bias from missing kinship excluded an entire branch of maternal relatives from the second generation across all simulated pedigrees. In reality, the unidentified kinship does not happen in all families in the data, and not all missed kinship occur on a maternal branch, which both cause less bias than demonstrated in our simulations. Simultaneously, unidentified kinship can exist in all the generations involved, and missing information in any generations apart from the second generation leads to less biased relatedness matrices. Although the bias analysis showed that the estimates of *mt*^2^ diminished about 40% when missing one piece of information to build the intact family tree, the impact of such misspecification is much less profound. The same discrepancy exists between the simulation of unidentified mtDNA mutation and the occurrence of these mutations in real data. However, we find that non-identified mtDNA mutation is linked with a 35% decrease of the *mt*^2^ estimates. Furthermore, we find that both unidentified kinship and unidentified mtDNA mutation leads to a strong overestimation of *j*^2^ estimates. The similarity between the related matrices of *a*^2^ and *j*^2^ are the primary force driving the overestimation. Based on these results, we recommend users of the current model to be more cautious about interpreting the results when the *j*^2^ estimate surpasses the *mt*^2^ estimate in the model fitting results, as the pattern could be a strong indication of incorrectly specified relatedness matrices.

The robustness of our model is further evidenced by its performance under the null model, when no mtDNA effects are present. Specifically, only a small proportion of mtDNA estimates—19.20%—deviated from zero by more than 0.01, underscoring the model’s capacity to discern true mtDNA variance from noise. Therefore, it is very unlikely to misattribute other sources of variance to mtDNA, if the mtDNA effects account for more than 2% of the total phenotypic variance. Datasets that have records of extended pedigrees typically encompass hundreds of thousands of individuals. For example, the Utah Population Database (UPB Smith et al., 2022; Bean, May, & Skolnick, 1978) has over 11 million people nested within pedigrees, crowd-sourced pedigrees hosted on Geni.com contain over 86 million people (including a single pedigree with 13 million people, Kaplanis et al., 2018) in publicly available pedigrees, and the Norwegian Mother, Father and Child Cohort Study (MoBa Magnus et al., 2016) has at least 95,000 families, sizes that are more than sufficient to detect mtDNA effects. The low false positive rate can ease researchers and funding agencies’ anxiety about spending efforts and funding on a spurious research direction. On the other side of the coin, the usage of current models will provide reliable insights on revealing the role of mtDNA in various phenotypes and effectively guide the future direction.

One limitation of the power analysis is that we only investigated continuous traits. Abundant evidence in family designs has suggested dichotomous traits will lead to a significant drop in bias(M. C. Neale, Eaves, & Kendler, 1994). We also anticipate the power for mtDNA estimation will decrease when applying the model to a binary outcome. Based on our results from a continuous outcome, large pedigree datasets with more than 20,000 individuals should be sufficient to detect a mtDNA effect of no less than 1%. However, an accurate power curve needs a similar simulation design to the current study so that it can be precisely derived.

One other notable limitation of the current study is the use of fixed-generation pedigrees for each condition in the simulations. Specifically, each group consisting of 20,000 simulated individuals has the same number of generations, though the number of offspring per mate varies among different mates following a Poisson distribution. The current approach offers advantages in power calculation because we can obtain an accurate unit non-centrality parameter for all sample sizes under the same conditions (Dominicus, Skrondal, Gjessing, Pedersen, & Palmgren, 2006; Satorra & Saris, 1985). However, real-life data more closely resembles a combination of pedigrees with varying depth. The impact of pedigree structure will qualitatively hold true, but the sample sizes to reach a certain threshold of power will be different.

We did not craft simulations for all possible violation assumptions, focusing instead on those we deemed most critical for this model—unidentified kinship and mtDNA mutations. We did not include other potential misspecifications that also bias the relatedness matrices. These potential misspecifications include treating twins and half-siblings as full siblings, as well as structural issues like endogamy (marriage within a specific community) and pedigree collapse (where ancestors appear multiple times in a pedigree). These scenarios are less common and generally result in only modest alterations to the relatedness matrices, thereby introducing less substantial bias compared to the primary issues we addressed. For researchers concerned that these less common scenarios might substantially affect their particular sample results, our (BGmisc Garrison et al., 2024), provides tools designed to analyze such biases, including specialized functionality for twinning and pedigree collapse. (See Hunter et al., 2024, for additional discussion on these tools.)

The model provides the field with a robust tool to explore the contribution of mtDNA on human complex traits. Our team has applied the model to longevity and Alzheimer’s disease, and some positive indication of mtDNA effects showed up in both trait (). Besides empirical application, a few future model modifications can be easily adapted to the current model. First, similar to other covariance structure models in behavioral genetics, this model can be extended to multivariate analysis, allowing for the estimation of shared mtDNA effects between two or more traits. Second, mtDNA effects can be incorporated into other extended family biometrical models as an additional source of phenotypic variance, given sufficiently separated maternal and paternal relatives. Third, with the emergence of genotyped family databases, modeling the influence of measured mtDNA haplogroups on top of the additive genetic effects and environmental effects is highly feasible and could further illustrate the biological effects of mtDNA on complex human traits.

## Conclusion

In the current study, we proposed and evaluated an extended pedigree model to quantify the influence of mtDNA on human behavior, controlling for influences from other genetic and environmental sources. The statistical power and bias analysis of the mtDNA effect estimate suggests that the model is sufficiently robust as a pioneering qualitative and quantitative exploration of the potential impact of mtDNA. The model is a powerful enough tool to detect even small mtDNA effects (5% of total variance) with a medium sample size (around 5,000). Nevertheless, we found that pedigree structures have a moderate influence on the statistical power of mtDNA effect estimation. Specifically, pedigrees with more generations typically increase power, while pedigrees with more offspring per mate typically decrease power, given the same total number of individuals involved. In terms of estimation bias, not having accurate kinship information and not accounting for mtDNA mutation underestimate the mtDNA variance by 35% to 40% under extremely misspecified scenarios. In addition, the mtDNA variance estimate has a small false positive rate. These findings substantiate the model’s value in examining the role of mtDNA in human behavior, and open up numerous opportunities for designing novel biometrical models and investigating new research questions in behavior genetics.

## Footnotes

1 Although these parameters are in the formats of squared coefficients, we do not impose a lower bound of 0 when estimating them in OpenMx.

